# Predicting future drinking among young adults: using ensemble machine-learning to combine MRI with psychometrics and behaviour

**DOI:** 10.1101/2020.03.03.974931

**Authors:** Martine M. Groefsema, Maartje Luijten, Rutger C.M.E. Engels, Guillaume Sescousse, Lee Jollans

## Abstract

**Background:** While most research into predictors of problematic alcohol use has focused on adolescence, young adults are also at elevated risk, and differ from adolescents and adults in terms of exposure to alcohol and neurodevelopment. Here we examined predictors of alcohol use among young adults at a 1-year follow-up using a broad predictive modelling approach.

**Methods:** Data in four modalities were included from 128 men aged between 18 and 25 years; functional MRI regions-of-interest from 1) a beer-incentive delay task, and 2) a social alcohol cue-exposure task, 3) grey matter data, and 4) non-neuroimaging data (i.e. psychometric and behavioural). These modalities were combined into an ensemble model to predict follow-up Alcohol Use Disorder Identification (AUDIT) scores, and were tested separately for their contribution. To reveal specificity for the prediction of future AUDIT scores, the same analyses were carried out for current AUDIT score.

**Results:** The ensemble resulted in a more accurate estimation of follow-up AUDIT score than any single modality. Only removal of the social alcohol cue-exposure task and of the non-neuroimaging data significantly worsened predictions. Reporting to need a drink in the morning to start the day was the strongest unique predictor of future drinking along with anterior cingulate cortex and cerebellar activity.

**Conclusions:** Alcohol-related task fMRI activity is a valuable predictor for future drinking among young adults alongside non-neuroimaging variables. Multi-modal prediction models best predict future drinking among young adults and may play an important part in the move towards individualized treatment and prevention efforts.

## Introduction

Alcohol consumption is highly prevalent during young adulthood, with 85% of young adults between 19-28 years reporting having tried alcohol, 31% having engaged in binge-drinking at least once in the last two weeks and around 4% reporting drinking on a daily basis in the last 30 days (1). This heavy drinking is accompanied by broad health risks and large societal costs (2–4). Luckily, many individuals ‘mature out’ of heavy drinking and decrease their alcohol intake when getting older (5–8). Currently, however, it remains difficult to identify who will reduce their alcohol consumption, who will continue to drink heavily, and who might even develop an alcohol dependence after drinking heavily.

Using machine learning, it is possible to predict complex clinical and behavioural outcomes by examining many variables (also referred to as “features”) simultaneously (9–11). Moreover, by considering predictors from a range of different modalities, the accuracy of predictions improves (e.g. (12)). Problematic alcohol use is a complex, multi-faceted behaviour related to biological, psychological and social factors (13–15), and is likely to be best explained by considering multiple predictors from multiple domains. A previous study that used such a multi-domain approach aimed to predict binge drinking in a large sample of adolescents (16). While the domains of personality and life history accounted for the majority of explained variance, other data domains also contributed unique variance (16). The results of two other machine learning studies revealed a variety of (f)MRI predictors for the initiation of alcohol use by the age of 18 (17), as well as non-neuroimaging predictors for the frequency of alcohol use among drinking and non-drinking adolescents (18).

Whereas these studies identified important risk factors associated with the initiation and continuation of drinking in young adolescents (i.e. an average age of 13), it is worthwhile to widen the scope of research into predictors of alcohol use to include early adulthood. First, prevalence rates of heavy drinking are high in this population, whilst research in this age group is scarce. Second, this population differs from adolescent and older adult samples in that alcohol use has already been initiated, but the likelihood of alcohol-related brain changes remains low. More specifically, it has been found that factors that predict alcohol use in adolescence are not necessarily predictive in young adults (19–21) and this may be related to adolescents undergoing continuing profound neurobiological changes (22). Moreover, the importance of neurobiological versus psychometric and behavioural risk factors may be different before and after alcohol use initiation, and therefore in adolescents versus young adults. For example, alcohol expectancies change over time and specifically after initiation (23, 24). Examining a wide range of possible predictors in young adults will help to identify risk factors that could contribute to intervention efforts specifically targeting this population.

The current study aims to identify predictors of heavy drinking in young adults who have already started drinking, by using a broad predictive modelling approach. In line with recommendations regarding the development of prediction models in psychiatry (25), our findings are intended to inform the design of future studies aimed at creating tools to identify at-risk individuals. To pursue this aim, we used an existing dataset that included a variety of potential neuroimaging predictors (i.e. grey matter volume and functional data from two different tasks including brain responses to alcohol cues within social settings, as well as brain responses to the anticipation and receipt of sips of beer (26, 27)) and psychometric predictors (i.e. non-neuroimaging data including self-report questionnaires and approach biases). Even though this dataset was originally not designed for predictive modelling, we were able to utilize this longitudinal dataset (including variables examined in previous publications, see Supplementary Table 1), to conduct an initial exploration of factors predicting problematic drinking in early adulthood.

## Materials and Methods

### Participants and procedure

The data used in this study were part of a larger project (see (27, 28) for a detailed description of the procedure and experimental tasks). At baseline, during two behavioural sessions and one fMRI session, a variety of measures were collected, using questionnaires or experimental task designs. One year later, data on alcohol use were obtained through self-report. Participants were men with varying levels of alcohol use (Table 2 for summary statistics). Differences between participants who did not complete follow-up measures (*n* = 19) and those who did (*n* = 128) did not survive multiple comparison (see Supplementary Figure 1 and Supplementary Table 2).

**Table 1.**
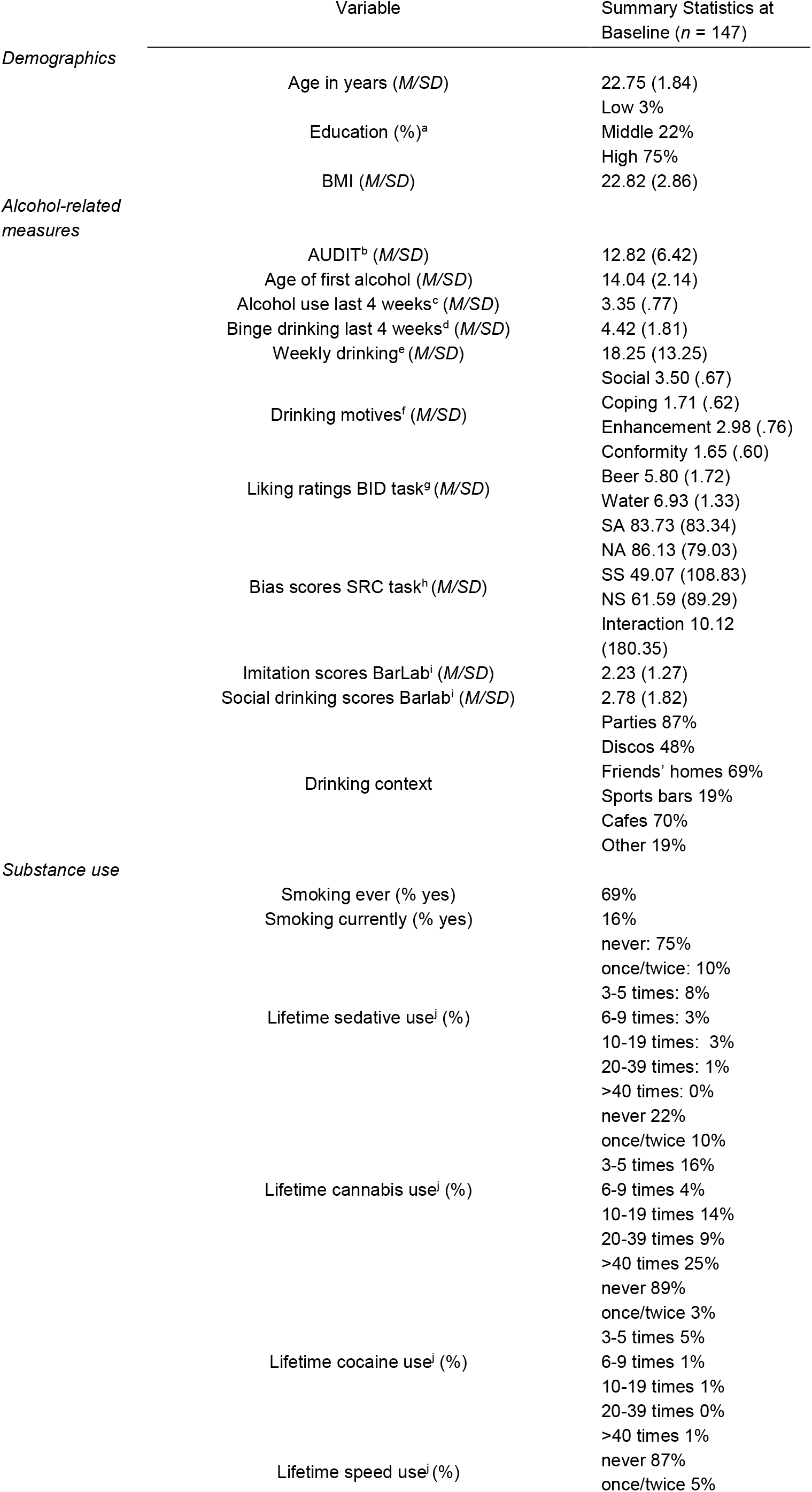
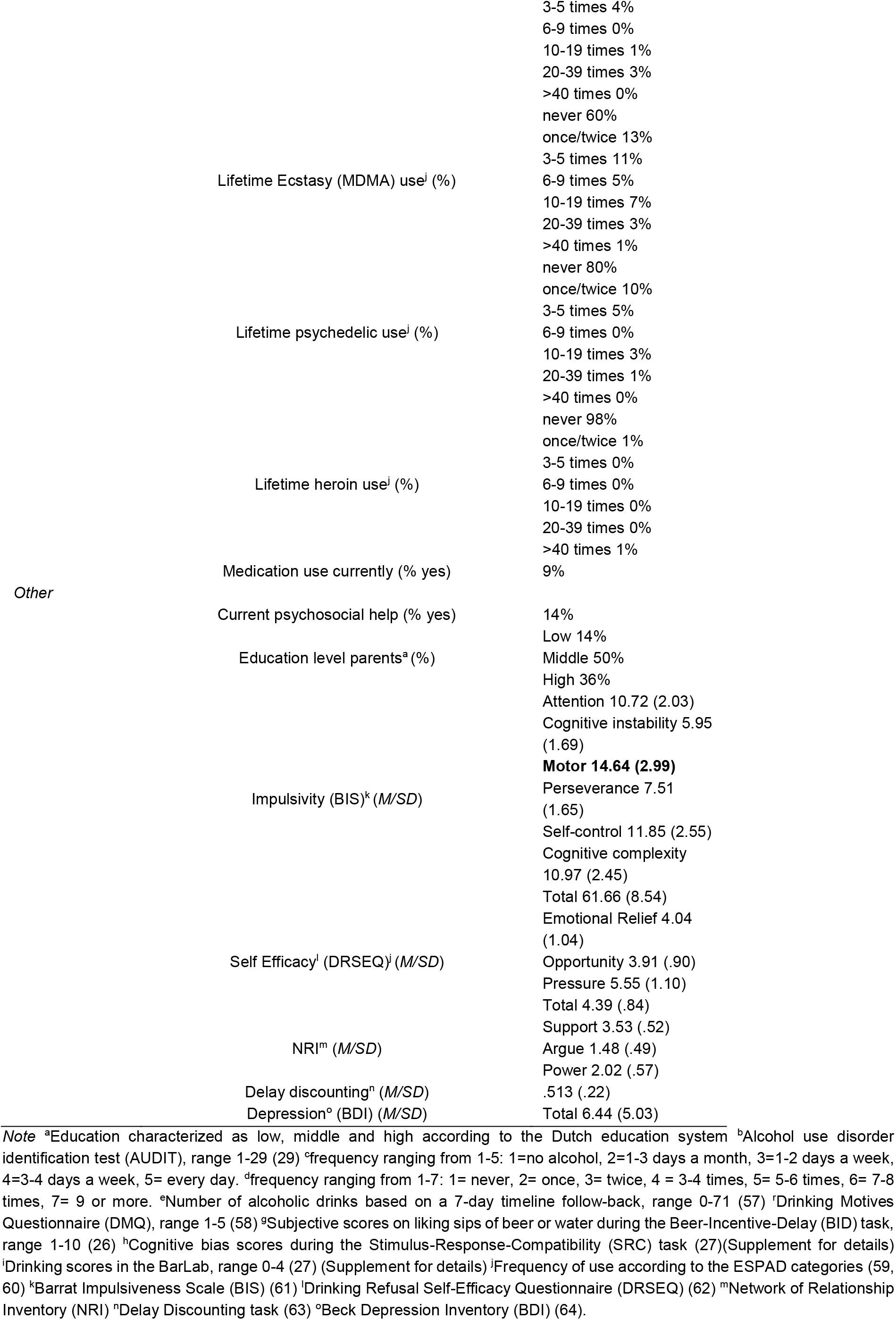
Sample descriptives of psychometric variables

### Measures

#### Outcome variable

Alcohol use was measured using the AUDIT questionnaire (29). Our sample (*n* = 128) had a mean AUDIT score of 12.82 (*SD* = 6.52) at baseline, and of 12.26 (*SD* = 5.78) at follow-up.

#### Input variables

##### Psychometric (non-neuroimaging)

The summary of demographic and cognitive variables (i.e. total scores or subscales) is reported in Table 1. For several questionnaires, single-item responses were included alongside summary values (i.e. for the AUDIT, weekly drinking, DMQ, BIS, NRI, DRSEQ, and BDI), resulting in 197 total features. For the AUDIT, the summary baseline score was included as a covariate in all analyses predicting follow-up AUDIT score, and lifetime heroin use was not included due to the low variability in the variable.

##### (f)MRI

During the MRI session, structural and functional data during two tasks were collected. One task was a passive cue-exposure task with four stimuli conditions consisting of alcohol or soda pictures, in a social or non-social setting (SACE) (27) for which the contrasts alcohol>soda, social>non-social, and ((social alcohol>social soda)>(non-social alcohol>non-social soda)) were included. The other task was a Beer-Incentive-Delay task with sips of beer or water as reinforcers (BID) (26), for which the contrasts beer>baseline, water>baseline, and beer>water for anticipation, outcome notification and delivery) of the task were included. See supplement for details of the pre-processing pipeline. Using a functionally defined atlas with 278 regions of interest (ROIs) (30), we extracted individual beta values for grey matter (278 features), and each of the SACE and BID contrasts (834 SACE features and 2502 BID features).

##### Analyses

To determine whether any identified predictors were specific to future drinking, we also performed the same analyses on current drinking. All analyses were carried out in MATLAB 2018b (scripts available at https://github.com/ljollans/multimodal_AUDIT_prediction). Given the relatively small sample size and large number of features for the BID and SACE task, standard multiple regression paired with bootstrap aggregation (bagging) was selected for all analyses (39), based on a previous empirical evaluation of the utility of various linear regression methods for prediction with neuroimaging data (31).

Models were built independently for the following four data domains: (1) fMRI ROIs for BID, (2) fMRI ROIs for SACE, (3) grey matter volume ROIs, and (4) non-neuroimaging variables. For each modality, a prediction model was built (“single-modality” model), generating an outcome prediction for each participant. Covariates included in all single-modality analyses were age and total AUDIT score at baseline. The four single-modality outcome predictions per participant were then treated as new predictors and entered into a further prediction model (i.e. an ensemble; (32)), with only those four variables (Figure 1). Outcome estimates from the ensemble reflect the extent to which combining data modalities results in a better prediction than using only data from one modality. To directly test this and measure the unique variance accounted for by each modality, the ensemble was also tested with each of the four data modalities excluded.

The entire analysis was carried out within a nested 5-fold cross-validation (CV) framework, that is, with an additional layer of CV within the training set of the main CV framework (33). The single-modality models were constructed in a nested training set (80% of the training set or ~64% of the whole dataset) and applied to the nested test set (20% of the training set or ~16% of the whole dataset). In the main training set (~80% of the data), the nested test set predictions were used to build the ensemble. This ensemble model was then applied to the outer test set (20% of observations) to generate a final outcome prediction for each observation. The entire analysis was repeated 10 times with 10 different CV assignments.

To determine whether the modality-specific models and the ensemble predicted the outcome significantly better than chance, all analyses were repeated using a randomly permuted outcome instead of the true outcome for each participant (“null” model) to establish an empirical significance threshold. The primary metric for estimating model fit was root mean squared error (RMSE) (33). Statistical significance was evaluated using students’ t-tests to compare RMSE values to RMSE values from the null model. For the single-modality models this was done using RMSE in the nested test sets (5 values for each iteration), and for the ensemble this was done using RMSE for the main test sets of all analysis iterations (one value for each iteration). For the single-modality models, predictors that significantly contributed to the model were also determined based on the null model. More specifically, those predictors for which the magnitude of the regression weights exceeded the 95^th^ percentile of the distribution seen in the null model were taken to contribute to the regression model above chance level (i.e. “passing the significance threshold”). Since the single-modality models generated in the nested CV framework and ensemble models generated in the main CV framework were evaluated, the significance threshold was based on the following levels of examination: (1) Average regression weights for each main CV iteration (5 sets of values per analysis iteration), (2) average regression weights for each analysis iteration (1 value per analysis iteration) (3) overall average regression weight. Subsequently, only predictors that passed the significance threshold (1) in at least one CV fold of every iteration, (2) in the majority of analysis iterations, and (3) when considering overall average regression weights, are reported. A full list of features and their performance for these metrics can be found in the Supplementary materials.

**Figure 1.**
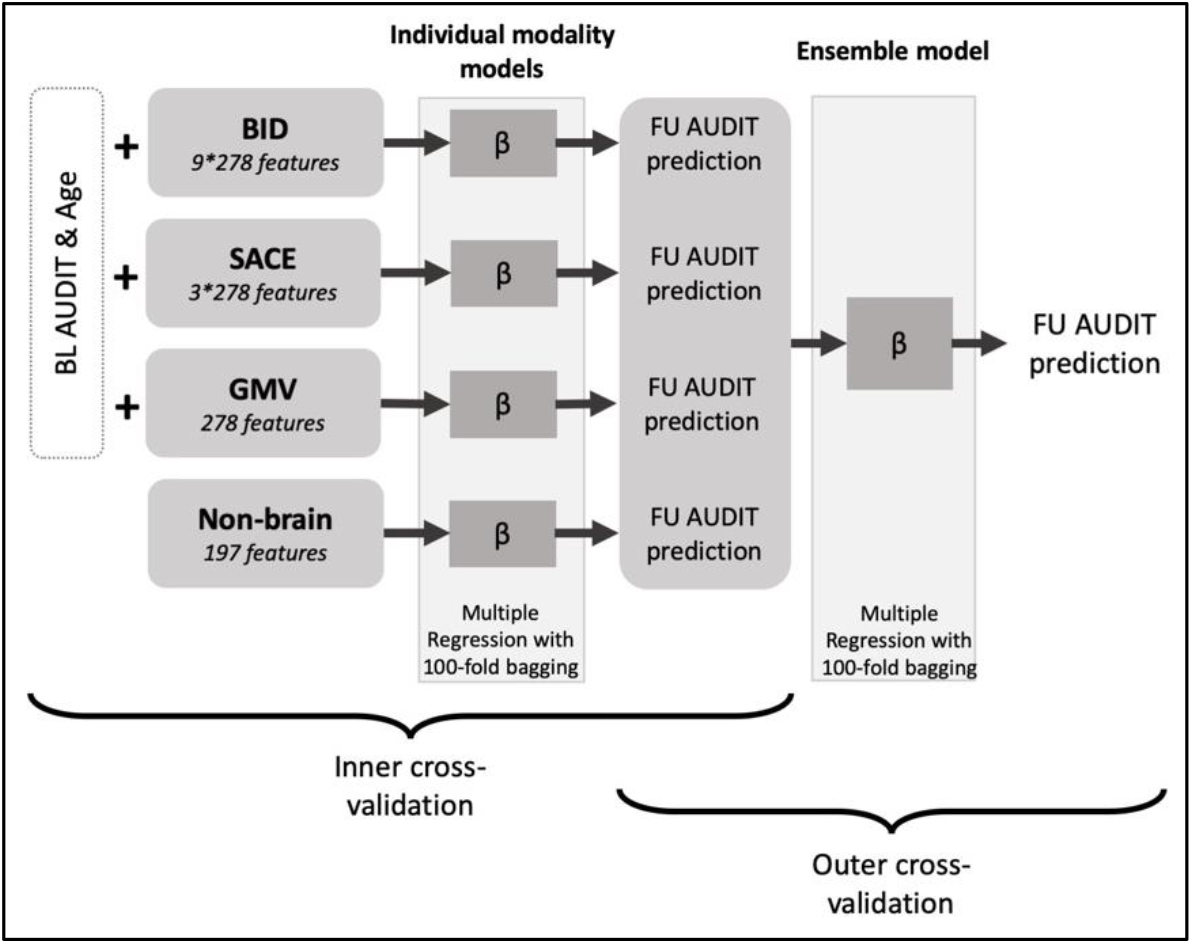
Representation of the analysis flow for ensemble learning models. BL: Baseline, FU: Follow-up (29), Beer-Incentive-Delay task (26), SACE: Social-Alcohol-Cue-Exposure task (27), GMV: Grey Matter Volume.

## Results

The ensemble model significantly predicted follow-up AUDIT score (p_RMSE_=6.38e-19, RMSE_avg_=0.6544, RMSE_min_=0.6299, RMSE_max_=0.6948). Pearson correlations indicate that between 51% and 60% of the variance in follow-up AUDIT score could be explained by the model (r_mean_=.7543, r_min_=.7190, r_max_=.7759,). All single-modality models were significantly better than chance (Table 2), but the ensemble was better than all single-modality models (p_BID_=1.5e-17, p_SACE_=6.5e-17, p_GMV_=5.7e-33,p_Psychometric_=1.7e-12).

**Table 2.** Single-modality and ensemble model fit

|  | BID | SACE | GMV | Psychometric |
| --- | --- | --- | --- | --- |
| Single-modality RMSE | .7810 | .7833 | .9172 | .7612 |
| (*p* compared to null) | (*p*=9.4e-51) | (*p*=3.1e-56) | (*p*=2.1e-41) | (*p*=6.5e-45) |
| Individual model *r*<sup>2</sup> | .3918 | .4137 | .3282 | .4353 |
| (*p* compared to null) | (*p*=2.4e-57) | (*p*=3.6e-61) | (*p*=4.6e-50) | (*p*=3.3e-54) |
| Increase in ensemble RMSE when<br>excluding this modality<br>(*p* compared to full ensemble) | 3.76%<br>(*p*=.0462) | 6.62%<br>(*p*=.0010)* | 2.50%<br>(*p*=.0978) | 13.09%<br>(*p*=5.9e-8)* |

\**p*<.0125 **Figure 1.** Representation of the analysis flow for ensemble learning models. BL: Baseline, FU: Follow-up (29), Beer-Incentive-Delay task (26), SACE: Social-Alcohol-Cue-Exposure task (27), GMV: Grey Matter Volume.

When comparing the ensemble model predicting follow-up AUDIT scores with all four modalities to a version of the ensemble where one modality was removed, removal of the psychometric and of the SACE task modality each resulted in a significant reduction in model performance after multiple comparison correction (Table 2). For baseline AUDIT scores, only the removal of the psychometric modality significantly reduced model fit, suggesting the SACE task modality is more useful for predicting follow-up AUDIT scores than for explaining baseline AUDIT scores (Supplementary Table 3).

### 1.0 Non-neuroimaging predictors

Current smoking and a higher score on AUDIT item 6 (“How often during the last year have you needed a first drink in the morning to get yourself going after a heavy drinking session?”) predicted higher AUDIT scores at follow-up.

Since current smoking was also predictive for baseline AUDIT scores, AUDIT item 6 was the only unique predictor of follow-up AUDIT scores (Supplementary Table 4 for unique predictors for baseline AUDIT scores)

### 2.0 Neuroimaging predictors

#### 2.1 SACE

##### 2.1.1 Alcohol>Soda

During the presentation of alcohol>soda pictures, higher activity in the medulla and left inferior parietal lobule (BA40), and lower activity in the lateral cerebellum and left ACC (BA24) predicted higher follow-up AUDIT scores.

The results for the prediction of baseline AUDIT scores showed that higher activity in the medulla was also predictive of higher baseline AUDIT scores, making activity in the inferior parietal lobule, cerebellum and ACC unique predictors for follow-up AUDIT scores (Figure 2).

##### 2.1.2 Social>Non-social

During the presentation of social>non-social pictures, lower activity in the left ACC and left cerebellum predicted higher follow-up AUDIT scores.

In contrast, the results for the prediction of baseline AUDIT scores showed that higher activity in the cerebellum predicted higher baseline AUDIT scores, with no effect for the same ACC region (Figure 2).

##### 2.1.3 Interaction ((social alcohol>social soda)>(non-social alcohol>non-social soda))

During the interaction contrast, higher activity in the right anterior PFC (BA10), and lower activity in the left occipital lobe (BA18), left thalamus, and cerebellum predicted higher follow-up AUDIT scores.

The results for the prediction of baseline AUDIT scores showed that lower activity in the same occipital ROI was also predictive for higher baseline AUDIT scores, making activity in the anterior PFC, thalamus, and cerebellum unique predictors for follow-up AUDIT scores (Figure 2).

**Figure 2.**
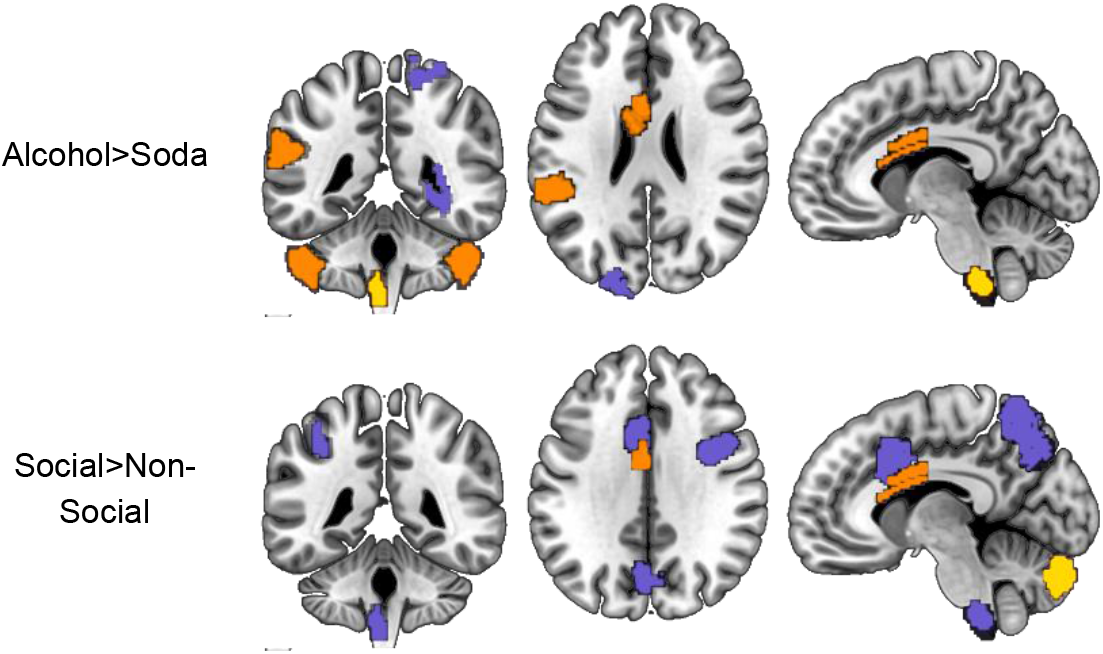
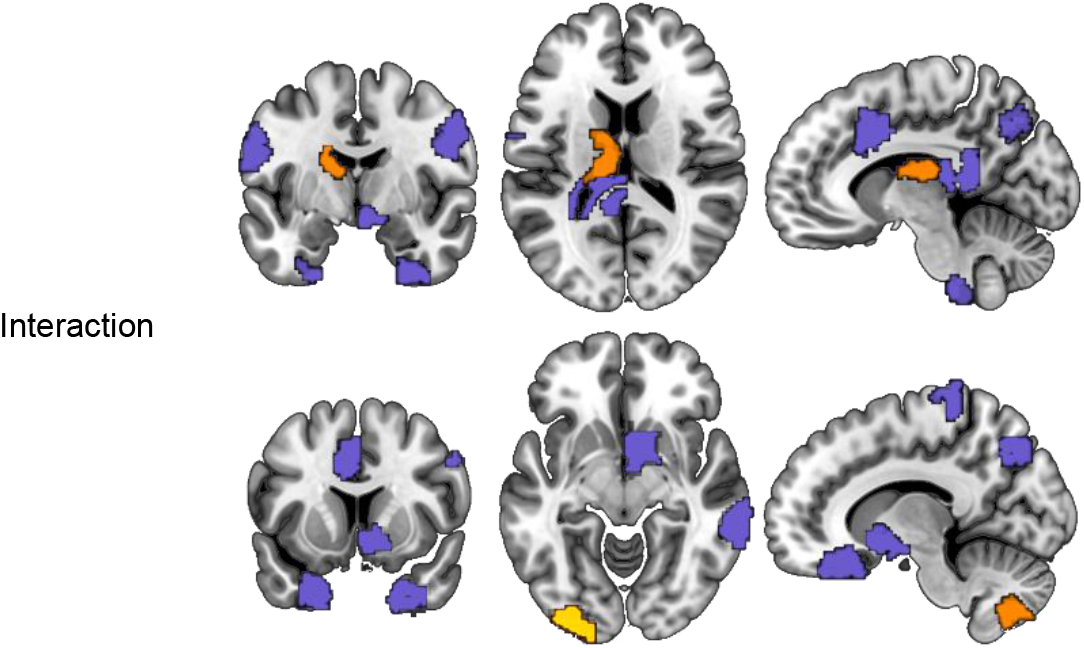
ROIs during the social alcohol cue-exposure task that were associated with the prediction of baseline AUDIT score (purple), follow-up AUDIT score (orange), or both (yellow).

#### 2.2 BID

For conciseness of the manuscript only the beer>water contrast for the three phases of the task is described here. A full description of all predictors for baseline as well as follow-up AUDIT scores in all contrasts can be found in Supplementary Table 4.

##### 2.2.1 Anticipation

During the anticipation of beer>water, higher activity in the left anterior dlPFC (BA 10 and BA 47) and right caudate nucleus (BA 48), and lower activity in the left primary motor cortex (BA 4) predicted higher follow-up AUDIT scores. With all areas except for the primary motor cortex being significant for baseline AUDIT prediction, the primary motor cortex activity was uniquely predictive of follow-up AUDIT (Figure 3).

##### 2.2.2 Outcome notification

During the outcome notification of beer>water, higher activity in the following areas predicted higher follow-up AUDIT scores: the bilateral OFC (BA 11) extending into the left nucleus accumbens (BA49), hypothalamus and subthalamic nuclei (BA49), the inferior frontal gyrus (BA45), left posterior cingulate cortex (BA30) extending into the caudate and fornix and adjacent to the lateral ventricle, left vermis (BA 19), left inferior temporal gyrus (BA 20). Moreover, lower activity in the right cerebellum and left ACC (BA24) was predictive of higher follow-up AUDIT scores.

The results for the prediction of baseline AUDIT scores showed that lower activity in the OFC and accumbens predicted higher baseline AUDIT, the effect being in the opposite direction to the follow-up AUDIT prediction. Results for the cerebellum, inferior frontal gyrus, and caudate/fornix were in the same direction for the baseline AUDIT prediction as for the follow-up AUDIT prediction. Vermis, inferior temporal gyrus, posterior and ACC activity were thus unique predictors of follow-up AUDIT (Figure 3).

##### 2.2.3 Delivery

During the delivery of beer>water, higher activity in the right cerebellum, the right anterior PFC (BA10), right premotor cortex (BA 8), left occipital cortex (BA 19), and brainstem predicted higher follow-up AUDIT scores. Moreover, lower activity in the left temporal cortex (BA 20, 21, 38) predicted higher follow-up AUDIT scores (Figure 3).

Activity in the right anterior PFC and right cerebellum were also predictive of baseline AUDIT, making premotor, occipital, brainstem, and temporal activity unique predictors of follow-up AUDIT (Figure 3).

**Figure 3.**
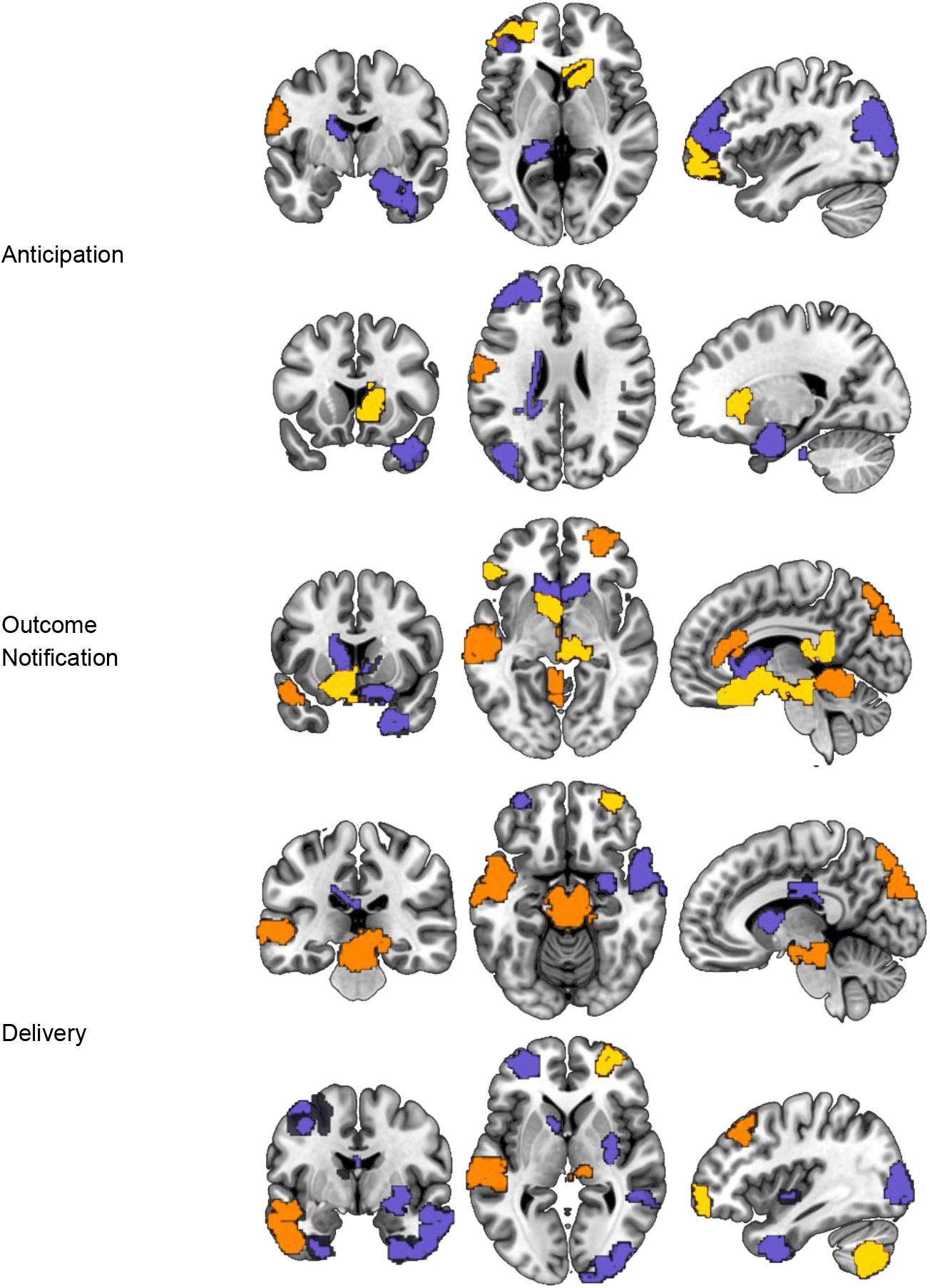
ROIs during the beer-incentive delay task that were associated with the prediction of baseline AUDIT score (purple), follow-up AUDIT score (orange), or both (yellow).

#### 2.3 GMV

Greater grey matter volume in the left insula (BA13), and lower grey matter volume in the right OFC (BA10) stretching into the anterior PFC (BA11) predicted higher follow-up AUDIT scores.

The results for the prediction of baseline AUDIT scores showed that more grey matter volume in the insula was also predictive for baseline AUDIT scores, making grey matter volume in the OFC and anterior PFC unique predictors for follow-up AUDIT scores (Figure 4).

**Figure 4.**
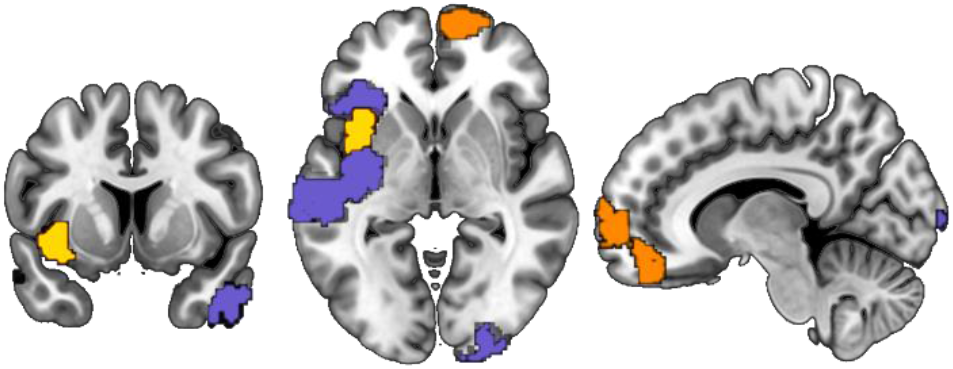
GMV ROIs during the beer-incentive delay task that were associated with the prediction of baseline AUDIT score (purple), follow-up AUDIT score (orange), or both (yellow).

## Discussion

Using a broad predictive modelling approach, the aim of this study was to identify variables that predict future heaviness of drinking in young adults, measured using AUDIT scores (29). The results revealed that an ensemble model, combining non-neuroimaging and neuroimaging data, predicted future drinking best. When testing all domains separately, they all performed better than chance, but only the removal of non-neuroimaging and cue-reactivity during the social cue-exposure task significantly increased error. Individuals reporting the need to drink in the morning was the strongest unique predictor. Thus, future drinking in young adults is most strongly predicted by disorder-like aspects of current drinking and BOLD responses to alcohol and social cues. Notably, neuroimaging data did not improve prediction of current alcohol use.

To the best of our knowledge, this is the first study to employ a data-driven machine learning approach to reveal variables that predict future alcohol use in young adult drinkers. While our results are in line with previous findings showing lower predictive value of neuroimaging compared to non-neuroimaging data for the prediction of heavy alcohol use over time (16, 34), we also demonstrated that the likelihood of heavy drinking is best explained by a combination of different assessment domains, implying complex and multi-faceted causal factors (16).

We identified that the need to drink in the morning, a variable not typically examined in similar studies (16–18), was the single strongest predictor of future drinking. This is therefore the first study to identify this simple self-report item as a possible target for risk assessment in young adults. Notably, this one item from the AUDIT questionnaire remained an important contributor to the prediction model despite controlling for overall AUDIT score. However, as previous similar studies have not tested this item specifically, replication is needed before use of this item as a screener in healthcare settings can be endorsed.

The results further demonstrate that BOLD responses to alcohol-related cues that include social environmental contexts contribute significantly to the prediction of future alcohol use, although inferential statistics reported in a previous publication did not find a relationship between social alcohol cue responses and actual drinking in social settings (27). Higher follow-up AUDIT scores were generally predicted by 1) lower activity in the ACC and cerebellum in response to alcohol (compared to soda) cues and in response to social (compared to non-social) cues, 2) higher activity in the inferior parietal lobule ((IPL) in response to alcohol (compared to soda) cues, 3) as well as higher activity in the PFC and lower activity in the thalamus and cerebellum in response to social alcohol (over social soda, compare to non-social alcohol over non-social soda) cues. These findings are not in line with the increased BOLD responses in reward-related brain areas that would be hypothesized based on the cue-reactivity literature (35–37). Instead, our findings seem to highlight the role of more cognitive control-related areas such as the ACC (38–40) during alcohol cue-reactivity as predictors of future use. Reduced activation in these areas is in line with dual-process theories (41) and the involvement of a fronto-parietal brain network in substance use. The current evidence is not sufficient to draw strong conclusions but supports increased attention to areas outside of the reward system in young adult substance use. In line with this argument, specific attention may be given to the role of the cerebellum. The cerebellum is a brain region that has consistently demonstrated volumetric (42–47) and functional abnormalities (48–51) in individuals with heavy alcohol consumption. To date, there have been few investigations of cerebellar function in young adults in relation to impulsive behaviour and substance use. However, reduced reward responses in binge-drinkers in the cerebellum have previously been observed (52), and our findings suggest that aberrant cerebellar function is associated with future drinking.

While the contribution to the multi-modal ensemble model was not statistically significant, BOLD responses during the anticipation and obtaining of sips of beer nevertheless predicted future AUDIT scores better than chance. Previous work using inferential statistics again found no association between BOLD responses during this task and different levels of problematic drinking (26). Without drawing strong conclusions, we want to highlight that higher activity in reward-related brain areas (OFC and ventral striatum) during anticipation was predictive of current and future drinking, suggesting that anticipation of beer rewards during this task is associated with real-life alcohol use. Moreover, lower activity in the ACC during outcome notification of beer compared to water predicted future drinking, but not current drinking. As noted above, low ACC activity during the cue-reactivity task was also a predictor of a higher follow-up AUDIT score, highlighting ACC responsiveness to alcohol stimuli as a risk factor for future heavy drinking.

Finally, while grey matter structure again predicted follow-up AUDIT scores better than chance, this modality predicted drinking with the lowest accuracy. Nevertheless, reduced grey matter volume in the insula and adjacent regions was associated with both current and future AUDIT scores, in line with a large body of work showing alterations in insular structure and function associated with substance use (53). We further found that reduced grey matter volume in the OFC and anterior PFC predicted future drinking only. Previous studies have found lower OFC volumes in heavy drinkers (54) and relapsers (55). Whether OFC volumetric changes are a pre-existing risk factor for alcohol use thus remains to be examined.

There are some important considerations to be noted regarding findings in this study. First, despite contributing the largest amount of explained variance to the prediction, only two of the non-neuroimaging variables reached the threshold for statistical significance. However, it is not necessarily the case that these two predictors contribute more to the AUDIT prediction than all other features. Since we examined findings at multiple stages of the cross-validation framework, the significance threshold was correspondingly very strict to minimize the chance of false positive findings. Furthermore, we did not exclude variables based on collinearity, and we included single-item values as well as the resulting scale summary values to maximize the information included in our dataset. Interaction effects or shared signal between features may thus have led to regression weights being distributed across features. Any individual feature sharing in such an effect may thus not have reached the significance threshold despite its inclusion contributing meaningfully to the model. As our primary aim was to quantify the overall contribution of different data modalities, further analyses to quantify the contribution of individual features or feature groups to the prediction was beyond the scope of this study. Second, despite including a large sample of participants for a neuroimaging study (*n* = 147 for current and *n* = 128 for future drinking), the sample is still relatively small in the context of machine learning. To mitigate this and enhance the reliability of our findings, we used multiple resampling techniques. With the field progressing towards larger sample sizes with more power (56), we encourage future studies to keep moving in this direction. Third, the implications of our findings are limited by the original study design. More specifically, only a relatively short time period elapsed between our baseline and follow-up measure, and our outcome measure consisted of self-report of heaviness of drinking (i.e. AUDIT) rather than a measure of real-time alcohol use.

In conclusion, here we used machine learning tools to gain insight into the relative contributions of predictors from different assessment domains for estimating future young adult alcohol use. We report four single-modality models reflecting brain structure, brain function, and non-neuroimaging variables including approach biases, life history, and personality factors, all of which were able to predict future alcohol use significantly above chance level. Furthermore, combining assessment domains into a single ensemble model significantly improved predictions. We can conclude that brain function and structure, as well as a wide array of self-report variables, can serve as valuable predictors for future heaviness of drinking among young adults. We found that the specific assessment of the urge to drink alcohol in the morning might be an important novel marker to identify future heavy drinkers. Based on our findings, we would further suggest a closer examination of the role of ACC and cerebellum activity and OFC grey matter volumes as potential risk factors associated with heavy drinking in young adults over a prolonged timeframe. Moreover, we illustrated that multi-modal prediction models can contribute valuable information to our understanding of the aetiology of heavy alcohol use and we encourage future studies to continue this line of research to eventually move towards individualized treatment and prevention efforts.

## Acknowledgments

Funding for this study was provided by the Behavioural Science Institute. The Behavioural Science Institute had no role in the study design, collection, analysis, or interpretation of the data, writing the manuscript, or the decision to submit the paper for publication. Using the same dataset, two papers have been published (26, 27).

