## Supplementary Material for "Predicting future drinking among young adults: using ensemble machine-learning to combine MRI with psychometrics and behaviour"

**Supplementary Table 1**. Overview of all collected data in the study

| WHEN | WHAT | HOW |
|---|---|---|
| **SCREENING** | | |
| | Alcohol use disorder | AUDIT |
| | Drinks last week | TLFB |
| | Gender | Male/female question |
| | Age | Open question |
| **MINI INTERVIEW (for the dependent group only)** | | |
| | Alcohol dependence diagnosis | MINI |
| | Other substance use information | MINI |
| **BEHAVIOURAL SESSION 1** | | |
| | Education | Multiple choice |
| | Age of first alcohol consumption | Open question |
| | Frequency of drinks in the last 4 weeks | Multiple choice |
| | Binge drinking episodes in last 4 weeks | Multiple choice |
| | Location of alcohol consumption | Multiple choice |
| | Drinking motives | DMQ-R |
| | Ever drank alcohol | Yes/No question |
| | Number of hours since last drink | Open question |
| | Ever smoked in life | Yes/No question |
| | Current smoker | Yes/No question |
| | Number of hours since last cigarette | Open question |
| | Smoking severity | FTND |
| | Ever used drugs (sleeping pills/cannabis/cocaine/ecstacy/amphetamine/hallucinogens/opiates), if yes how many times | Open question |
| | Impulsivity | BIS11 |
| | State-trait Anxiety | STAI |
| | Depression | BDI |
| | Drinking Urge | DAQ |
| | Imitation of alcohol use | Number of drinks consumed in Bar-lab |
| | Confederate liking ratings | 1-9 scales |
| | Anxiety before/during/after session | Multiple choice |
| | Suspicion checks about study goals | Open questions |
| **BEHAVIOURAL SESSION 2** | | |
| | Number of hours since last drink | Open question |
| | Number of hours since last cigarette | Open question |
| | Delay Discounting | Delay discounting task |
| | Drinking Urge | DAQ |
| | Imitation of alcohol use | Number of drinks consumed in Bar-lab |
| | Confederate liking ratings | 1-9 scales |
| | Anxiety before/during/after session | Multiple choice |
| | Suspicion checks about study goals | Open questions |

*Table S1. Overview of all collected data in the study (continued)*

| WHEN | WHAT | HOW |
|---|---|---|
| **FMRI SESSION** | | |
| | Number of hours since last drink | Open question |
| | Number of hours since last cigarette | Open question |
| | Height/Weight | Open question |
| | Drinking Urge | DAQ |
| | Drinking Self-Efficacy | DRSEQ |
| | Brain responses to social alcohol cues | Social Alcohol cue Reactivity task |
| | Brain responses to anticipating and receiving beer | Beer Incentive Delay task (current paper) |
| | Approach/Avoidance of (social) alcohol pictures | Stimulus-Response Compatibility task |
| **FOLLOW UP BASELINE** | | |
| | Alcohol use disorder | AUDIT |
| | Average number of drinks for each day of the week | Weekly drinking |
| | Drinking motives | DMQ-R |
| **FOLLOW UP WITH ECOLOGICAL MOMENTARY ASSESSMENT (14 DAYS)** | | |
| | Number of (non)alcohol units consumed day before | Open question |
| | Location of (non)alcohol consumption day before | Multiple choice |
| | People with whom (non)alcohol was consumed with day before | Multiple choice |

**Supplementary Table 2.** *Demographic, substance use, psychosocial and alcohol-specific characteristics of participants*

| | Variable | Summary Statistics of excluded participants due to missing follow-up AUDIT (*n* = 19) | Statistics (difference between included and excluded participants) |
|---|---|---|---|
| *Demographics* | | | |
| | Age in years (*M/SD*) | 22.43 (1.65) | *t*(145) = .820, *p* = .413 |
| | | Low 5% | |
| | Education (%)[a] | Middle 21% | $X^2(2)$ = .231, *p* = .891 |
| | | High 74% | |
| | BMI (*M/SD*) | 22.41 (2.23) | *t*(145) = .668, *p* = .505 |
| *Alcohol-related measures* | | | |
| | AUDIT[b] (M/SD) | 13.00 (6.13) | *t*(145) = .128, *p* = .898 |
| | Age of first alcohol (*M/SD*) | 14.26 (1.62) | *t*(145) = .468, *p* = .641 |
| | Alcohol use last 4 weeks[c] (*M/SD*) | 3.47 (.69) | *t*(145) = .722, *p* = .471 |
| | Binge drinking last 4 weeks[d] (*M/SD*) | 5.26 (1.91) | ***t*(145) = 2.191, *p* = .030** |
| | Weekly drinking[e] (*M/SD*) | 18.76 (11.52) | *t*(145) = .179, *p* = .859 |
| | Drinking motives[f] (*M/SD*) | Social 3.78 (.64) | ***t*(145) = 2.014, *p* = .046** |
| | | Coping 1.53 (.47) | *t*(145) = 1.341, *p* = .182 |
| | | Enhancement 3.13 (.68) | *t*(145) = .924, *p* = .357 |
| | | Conformity 1.71 (.42) | *t*(145) = .465, *p* = .642 |
| | Liking ratings BID task[g] (*M/SD*) | Beer 6.00 (1.37) | *t*(145) = 5.34, *p* = .594 |
| | | Water 6.42 (1.67) | *t*(145) = 1.805, *p* = .073 |
| | Bias scores SRC task[h] (*M/SD*) | SA 95.39 (78.45) | *t*(145) = .652, *p* = .515 |
| | | NA 69.57 (87.05) | *t*(145) = .978, *p* = .330 |
| | | SS 12.05 (102.31) | *t*(145) = 1.597, *p* = .112 |
| | | NS 34.23 (97.73) | *t*(145) = 1.436, *p* = .153 |
| | | Interaction 2.05 (1.02) | *t*(145)= .981, *p* = .328 |
| | Imitation scores BarLab[i] (*M/SD*) | 2.05 (1.02) | *t*(145)= .655, *p* = .514 |
| | Social drinking scores Barlab[i] (*M/SD*) | 3.42 (1.50) | *t*(145) = 1.630, *p* = .105 |
| | Drinking Context | Parties 89% | $X^2(1)$ = .112, *p* = .738 |
| | | Discos 47% | $X^2(1)$ = .008, *p* = .931 |
| | | Friends' homes 89% | **$X^2(1)$ = 4.377, *p* = .037** |
| | | Sports bars 5% | $X^2(1)$ = 2.689, *p* = .084 |
| | | Other 78% | $X^2(1)$ = .709, *p* = .590 |
| *Substance use* | | | |
| | Smoking ever (% yes) | 68% | $X^2(1)$ = .010, *p* = .922 |
| | Smoking currently (% yes) | 10% | $X^2(1)$ = .537, *p* = .364 |
| | Lifetime sedative use[j] (%) | never 79% | $X^2(5)$ = 2.008, *p* = .848 |
| | | once/twice 5% | |

| | | |
|---|---|---|
| | 3-5 times 11% | |
| | 6-9 times 0% | |
| | 10-19 times 5% | |
| | 20-39 times 0% | |
| | >40 times 0% | |
| | never 16% | |
| | once/twice 16% | |
| | 3-5 times 11% | |
| Lifetime cannabis use[j] (%) | 6-9 times 5% | $X^2(6) = 2.161$, $p = .904$ |
| | 10-19 times 10% | |
| | 20-39 times 10% | |
| | >40 times 32% | |
| | never 85% | |
| | once/twice 5% | |
| | 3-5 times 5% | |
| Lifetime cocaine use[j] (%) | 6-9 times 0% | $X^2(5) = 3.477$, $p = .627$ |
| | 10-19 times 5% | |
| | 20-39 times 0% | |
| | >40 times 0% | |
| | never 90% | |
| | once/twice 5% | |
| | 3-5 times 0% | |
| Lifetime speed use[j] (%) | 6-9 times 0% | $X^2(4) = 1.732$, $p = .785$ |
| | 10-19 times 0% | |
| | 20-39 times 5% | |
| | >40 times 0% | |
| | never 64% | |
| | once/twice 21% | |
| | 3-5 times 5% | |
| Lifetime Ecstasy (MDMA) use[j] (%) | 6-9 times 5% | $X^2(6) = 6.306$, $p = .390$ |
| | 10-19 times 0% | |
| | 20-39 times 0% | |
| | >40 times 5% | |
| | never 90% | |
| | once/twice 5% | |
| | 3-5 times 5% | |
| Lifetime LSD/paddo’s use[j] (%) | 6-9 times 0% | $X^2(4) = 1.644$, $p = .801$ |
| | 10-19 times 0% | |
| | 20-39 times 0% | |
| | >40 times 0% | |
| Lifetime heroin use[j] (%) | never 100% | $X^2(2) = .301$, $p = .860$ |

| | | | |
|---|---|---|---|
| | | once/twice 0% | |
| | | 3-5 times 0% | |
| | | 6-9 times 0% | |
| | | 10-19 times 0% | |
| | | 20-39 times 0% | |
| | | >40 times 0% | |
| | Medication use currently (% yes) | 0% | $X^2(1) = 2.117$, $p = .152$ |
| *Other* | | | |
| | Current psychosocial help (% yes) | 5% | $X^2(1) = 1.475$, $p = .199$ |
| | | Low 5% | |
| | Education level parents[a] (%) | Middle 58% | $X^2(2) = 1.367$, $p = .505$ |
| | | High 37% | |
| | Impulsivity (BIS)[k] (*M/SD*) | Attention 11.31 (2.56) | $t(145) = 1.353$, $p = .178$ |
| | | Cognitive instability 6.21 (1.98) | $t(145) = .709$, $p = .479$ |
| | | Motor 15.42 (1.86) | $t(145) = 1.209$, $p = .228$ |
| | | Perseverance 7.63 (1.77) | $t(145) = .342$, $p = .733$ |
| | | Self-control 12.52 (2.85) | $t(145) = 1.237$, $p = .218$ |
| | | Cognitive complexity 11.00 (2.72) | $t(145) = .039$, $p = .969$ |
| | | Total 64.10 (9.46) | $t(145) = 1.336$, $p =.184$ |
| | Self Efficacy[l] (DRSEQ)[j] (*M/SD*) | Emotional Relief 4.20 (1.10) | $t(145) = .711$, $p = .478$ |
| | | Opportunity 4.07 (.86) | $t(145) = .815$, $p = .416$ |
| | | Pressure 5.70 (1.14) | $t(145) = .621$, $p = .536$ |
| | | Total 4.55 (.83) | $t(145) = .867$, $p = .387$ |
| | NRI[m] (*M/SD*) | Support 3.37 (.56) | $t(145) = 1.434$, $p = .154$ |
| | | Argue 1.50 (.49) | $t(145) = .149$, $p = .882$ |
| | | Power 2.29 (.65) | **$t(145) = 2.222$, $p = .028$** |
| | Delay discounting[n] (*M/SD*) | .391 (.234) | **$t(145) = 2.567$, $p = .011$** |

*Note* [a]Education characterized as low, middle and high according to the Dutch education system [b]Alcohol use disorder identification test (AUDIT), range 1-29 (1) [c]frequency ranging from 1-5: 1=no alcohol, 2=1-3 days a month, 3=1-2 days a week, 4=3-4 days a week, 5= every day. [d]frequency ranging from 1-7: 1= never, 2= once, 3= twice, 4 = 3-4 times, 5= 5-6 times, 6= 7-8 times, 7= 9 or more. [e]Number of alcoholic drinks based on a 7-day timeline follow-back, range 0-71 (2) [f]Drinking Motives Questionnaire (DMQ), range 1-5 (3) [g]Subjective scores on liking sips of beer or water during the Beer-Incentive-Delay (BID) task, range 1-10 (4) [h]Cognitive bias scores during the Stimulus-Response-Compatibility (SRC) task (5)(Supplement for details) [i]Drinking scores in the BarLab, range 0-4 (5) (Supplement for details) [j]Frequency of use according to the ESPAD categories (6, 7) [k]Barrat Impulsiveness Scale (BIS) (8) [l]Drinking Refusal Self-Efficacy Questionnaire (DRSEQ) (9) [m]Network of Relationship Inventory (NRI) [n]Delay Discounting task (10) [o]Beck Depression Inventory (BDI) (11).

**Supplementary Table 3.** Single-modality and ensemble model fit for baseline AUDIT scores

| | BID | SACE | GMV | Psychometric |
|---|---|---|---|---|
| Single-modality RMSE<br>(*p* compared to null) | 1.1454<br>(*p*=6.0e-4) | 1.1548<br>(*p*=1.3e-12) | 1.367<br>(*p*=1.8e-6) | 0.9447<br>(*p*=7.2e-38) |
| Individual model $r^2$<br>(*p* compared to null) | . 0077<br>(*p*=1.8e-8) | .0223<br>(*p*=4.0e-15) | . 0338<br>(*p*=3.6e-19) | . 2230<br>(*p*=6.4e-49) |
| Increase in ensemble RMSE when excluding this modality<br>(*p* compared to full ensemble) | -1.71%<br>(*p*=.4490) | 2.29%<br>(*p*=.1818) | 1.44%<br>(*p*=.4010) | 65.22%<br>(*p*=1.5e-14)* |

*$p$<.0125

*Table S4. Predictors for Follow-Up and baseline prediction of AUDIT*

| Task | Contrast | | Follow-up Prediction | | | Baseline Prediction | | |
|---|---|---|---|---|---|---|---|---|
| Task | Contrast | | Shen Mask | Hemisphere | Beta weight | Shen Mask | Hemisphere | Beta weight |
| SACE | | | | | | | | |
| | | Alcohol>Soda | | | | | | |
| | | | **272_No_BA_region** | **-** | **0.040301873** | 105_R_BA6_2 | R | -0.031804272 |
| | | | 245_L_BA40_3 | L | 0.023742149 | 153_L_BA18_2 | L | -0.029190911 |
| | | | 159_No_BA_region | - | -0.028838425 | 179_L_BA47_1 | L | 0.029567214 |
| | | | 66_R_BA20_3 | R | -0.027184733 | **272_No_BA_region** | **-** | **0.085927676** |
| | | | 176_L_BA24_2 | L | -0.024824482 | 29_R_BA7_6 | R | 0.044614138 |
| | | | | | | 42_R_BA54_2 | R | 0.032166899 |
| | | | | | | 77_R_BA54_1 | R | -0.03476863 |
| | | | | | | 80_R_BA7_4 | R | 0.040903774 |
| | | Social>Nonsocial | | | | | | |
| | | | **256_No_BA_region** | **-** | **-0.025369967** | 116_R_BA6_6 | R | -0.028778514 |
| | | | 176_L_BA24_2 | L | -0.022075276 | 146_L_BA7_8 | L | -0.027156312 |
| | | | | | | 150_No_BA_region | - | 0.056104493 |
| | | | | | | 16_R_BA7_1 | R | -0.058105812 |
| | | | | | | 160_L_BA7_1 | L | 0.043437132 |
| | | | | | | 173_L_BA32_4 | L | -0.032169487 |
| | | | | | | 194_L_BA39_5 | L | 0.087558697 |
| | | | | | | 249_L_BA7_9 | L | -0.02727052 |
| | | | | | | **256_No_BA_region** | **-** | **0.065406702** |
| | | | | | | 257_L_BA18_1 | L | 0.031361111 |
| | | | | | | 272_No_BA_region | - | -0.072738141 |
| | | | | | | 6_R_BA10_4 | R | 0.044625982 |
| | | | | | | 63_R_BA8_6 | R | -0.051909154 |
| | | | | | | 85_R_BA6_5 | R | -0.040928253 |
| | | Interaction | | | | | | |
| | | | 6_R_BA10_4 | R | 0.021764506 | 114_R_BA55_1 | R | -0.053727855 |
| | | | **257_L_BA18_1** | **L** | **-0.03640035** | 154_L_BA4_1 | L | 0.03595173 |
| | | | 260_L_BA48_1 | L | -0.026800372 | 16_R_BA7_1 | R | 0.03222258 |
| | | | 76_No_BA_region | - | -0.024486113 | 173_L_BA32_4 | L | -0.038406156 |
| | | | | | | 195_L_BA40_1 | L | 0.031004542 |
| | | | | | | 204_L_BA30_1 | L | 0.027184315 |
| | | | | | | 22_R_BA11_3 | R | -0.058144912 |
| | | | | | | 233_L_BA50_1 | L | 0.036958843 |
| | | | | | | 249_L_BA7_9 | L | 0.031176383 |
| | | | | | | **257_L_BA18_1** | **L** | **-0.037815318** |
| | | | | | | 265_L_BA38_3 | L | 0.032603434 |
| | | | | | | 272_No_BA_region | - | -0.04248748 |
| | | | | | | 32_R_BA1_1 | R | -0.038841357 |
| | | | | | | 33_R_BA21_4 | R | -0.039098292 |
| | | | | | | 34_R_BA6_8 | R | 0.036080723 |

| | | | | | | | | |
|---|---|---|---|---|---|---|---|---|
| | | | | | | 70_R_BA38_1 | R | 0.035214278 |
| BID | | | | | | | | |
| | Anticipation phase | | | | | | | |
| | | Beer>Baseline | | | | | | |
| | | | 35_R_BA37_8 | R | -0.011347086 | 10_R_BA21_2 | R | -0.01193203 |
| | | | | | | 103_R_BA47_4 | R | 0.010048589 |
| | | | | | | 114_R_BA55_1 | R | 0.016617861 |
| | | | | | | 148_L_BA1_2 | L | 0.013738618 |
| | | | | | | 22_R_BA11_3 | R | -0.013473385 |
| | | | | | | 229_L_BA39_3 | L | 0.010758577 |
| | | | | | | 234_L_BA21_2 | L | -0.013259178 |
| | | | | | | 242_L_BA6_5 | L | 0.010459376 |
| | | | | | | 257_L_BA18_1 | L | -0.020413362 |
| | | Water>Baseline | | | | | | |
| | | | **126_No_BA_region** | **-** | **0.009164304** | 110_R_BA18_4 | R | 0.015881429 |
| | | | 173_L_BA32_4 | L | 0.011172339 | **126_No_BA_region** | **-** | **0.022487182** |
| | | | 180_L_BA32_1 | L | 0.009973149 | 257_L_BA18_1 | L | -0.02750998 |
| | | Beer>Water | | | | | | |
| | | | **179_L_BA47_1** | **L** | **0.015771535** | 112_R_BA20_1 | R | -0.017668314 |
| | | | **193_L_BA10_2** | **L** | **0.009256542** | 115_No_BA_region | - | -0.010145116 |
| | | | **25_R_BA48_1** | **R** | **0.012164176** | 123_R_BA38_2 | R | -0.035033761 |
| | | | 154_L_BA4_1 | L | -0.030225147 | 133_R_BA53_1 | R | -0.010395298 |
| | | | | | | 140_L_BA10_1 | L | 0.010711719 |
| | | | | | | **179_L_BA47_1** | **L** | **0.016327936** |
| | | | | | | 185_L_BA19_4 | L | 0.013839286 |
| | | | | | | **193_L_BA10_2** | **L** | **0.017236057** |
| | | | | | | 233_L_BA50_1 | L | 0.015202123 |
| | | | | | | **25_R_BA48_1** | **R** | **0.01858005** |
| | | | | | | 259_L_BA10_3 | L | 0.010912069 |
| | | | | | | 260_L_BA48_1 | L | 0.012806905 |
| | | | | | | 275_L_BA19_2 | L | 0.016077346 |
| | | | | | | 276_L_BA46_1 | L | 0.027009067 |
| | | | | | | 57_R_BA1_3 | R | -0.015380951 |
| | | | | | | 94_R_BA40_3 | R | -0.011955616 |
| | Outcome Notification | | | | | | | |
| | | Beer>Baseline | | | | | | |
| | | | 272_No_BA_region | - | 0.010158181 | 204_L_BA30_1 | L | 0.020199669 |
| | | | | | | 22_R_BA11_3 | R | -0.015268768 |
| | | Water>Baseline | | | | | | |
| | | | 223_L_BA19_9 | L | 0.008862958 | 107_R_BA9_5 | R | 0.012529196 |
| | | | 148_L_BA1_2 | L | 0.009250663 | 136_R_BA36_2 | R | 0.010936864 |
| | | | 165_L_BA18_8 | L | 0.009813495 | 152_L_BA23_2 | L | -0.01387106 |
| | | | 232_L_BA7_5 | L | 0.012363161 | 158_No_BA_region | - | 0.01865288 |
| | | | 39_R_BA19_5 | R | 0.014355088 | 16_R_BA7_1 | R | -0.012553118 |
| | | | **36_R_BA19_2** | **R** | **0.015352546** | 173_L_BA32_4 | L | -0.009723793 |

| | | | | | | | |
|---|---|---|---|---|---|---|---|
| | | 257_L_BA18_1 | L | -0.024145441 | 194_L_BA39_5 | L | 0.012505437 |
| | | **251_L_BA8_2** | **L** | **-0.014140336** | 248_L_BA9_3 | L | -0.010747647 |
| | | 129_R_BA10_6 | R | -0.010213014 | **251_L_BA8_2** | **L** | **-0.010671965** |
| | | | | | **36_R_BA19_2** | **R** | **-0.019158505** |
| | | | | | 4_R_BA1_2 | R | -0.014463235 |
| | | | | | 6_R_BA10_4 | R | 0.015769958 |
| | | | | | 78_R_BA47_3 | R | -0.019543225 |
| | Beer>Water | | | | | | |
| | | 204_L_BA30_1 | L | 0.010094383 | 100_R_BA40_2 | R | 0.011950036 |
| | | **22_R_BA11_3** | **R** | **0.013026181** | 110_R_BA18_4 | R | 0.009659846 |
| | | 175_L_BA19_10 | L | 0.01303595 | 112_R_BA20_1 | R | -0.013962783 |
| | | 198_L_BA20_1 | L | 0.016230551 | 126_No_BA_region | - | 0.013376615 |
| | | **151_L_BA49_2** | **L** | **0.018046265** | 13_No_BA_region | - | -0.017595045 |
| | | **170_L_BA11_2** | **L** | **0.018866061** | 139_R_BA6_7 | R | 0.014655441 |
| | | **233_L_BA50_1** | **L** | **0.020949916** | **151_L_BA49_2** | **L** | **-0.00928075** |
| | | **215_L_BA45_1** | **L** | **0.036437282** | **170_L_BA11_2** | **L** | **-0.042524522** |
| | | **66_R_BA20_3** | **R** | **-0.017156142** | **215_L_BA45_1** | **L** | **0.009955029** |
| | | 161_L_BA24_1 | L | -0.010916281 | **22_R_BA11_3** | **R** | **-0.032089909** |
| | | | | | **233_L_BA50_1** | **L** | **0.014965621** |
| | | | | | 25_R_BA48_1 | R | -0.018232736 |
| | | | | | 263_L_BA48_3 | L | -0.015247756 |
| | | | | | 273_L_BA48_2 | L | -0.010983663 |
| | | | | | 4_R_BA1_2 | R | 0.011734791 |
| | | | | | 6_R_BA10_4 | R | -0.012555677 |
| | | | | | **66_R_BA20_3** | **R** | **-0.014347442** |
| | | | | | 70_R_BA38_1 | R | 0.010715615 |
| | | | | | 74_R_BA47_1 | R | -0.012276403 |
| | | | | | 75_R_BA39_4 | R | 0.016096005 |
| Delivery | | | | | | | |
| | Beer>Baseline | | | | | | |
| | | | | | 272_No_BA_region | - | 0.033222571 |
| | Water>Baseline | | | | | | |
| | | 158_No_BA_region | **-** | -0.009920713 | 116_R_BA6_6 | R | -0.018227873 |
| | | 42_R_BA54_2 | R | -0.016909306 | 139_R_BA6_7 | R | -0.012664512 |
| | | 70_R_BA38_1 | R | -0.013789785 | 272_No_BA_region | - | 0.022858044 |
| | | | | | 28_R_BA4_1 | R | -0.013970208 |
| | | | | | 34_R_BA6_8 | R | -0.017304429 |
| | | | | | 70_R_BA38_1 | R | 0.009331427 |
| | Beer>Water | | | | | | |
| | | 76_No_BA_region | - | 0.014659192 | 10_R_BA21_2 | R | 0.013246259 |
| | | 223_L_BA19_9 | L | 0.00925614 | 120_R_BA18_7 | R | -0.014898575 |
| | | 58_R_BA50_2 | R | 0.010155588 | 121_R_BA18_6 | R | -0.017571307 |
| | | **6_R_BA10_4** | **R** | **0.01238789** | 122_R_BA37_10 | R | 0.013175647 |
| | | 144_L_BA55_1 | L | 0.012699303 | **126_No_BA_region** | **-** | **0.028390507** |

| | | | | | | |
|---|---|---|---|---|---|---|
| | **126_No_BA_region** | **-** | **0.014327508** | 140_L_BA10_1 | L | 0.011781666 |
| | 72_R_BA8_5 | R | 0.014659192 | 142_No_BA_region | - | -0.012024517 |
| | 232_L_BA7_5 | L | 0.015209754 | 152_L_BA23_2 | L | 0.010772799 |
| | 198_L_BA20_1 | L | -0.013872273 | 159_No_BA_region | - | -0.008207227 |
| | 210_L_BA21_4 | **L** | -0.012690334 | 193_L_BA10_2 | L | 0.023553666 |
| | 234_L_BA21_2 | **L** | -0.012034854 | 197_L_BA6_3 | L | 0.015629508 |
| | 244_L_BA38_1 | L | -0.010541888 | 203_L_BA6_6 | L | 0.012659086 |
| | | | | 26_R_BA20_2 | R | -0.021814024 |
| | | | | 265_L_BA38_3 | L | -0.010083499 |
| | | | | 273_L_BA48_2 | L | 0.011987059 |
| | | | | 47_R_BA23_1 | R | 0.010771693 |
| | | | | **6_R_BA10_4** | **R** | **0.021289696** |
| | | | | 70_R_BA38_1 | R | -0.056191491 |
| | | | | 75_R_BA39_4 | R | 0.013492614 |
| | | | | 89_R_BA38_3 | R | -0.011975584 |
| | | | | 98_R_BA49_2 | R | -0.020715617 |
| | | | | 10_R_BA21_2 | R | 0.013246259 |
| | | | | 120_R_BA18_7 | R | -0.014898575 |
| | | | | 121_R_BA18_6 | R | -0.017571307 |
| | | | | 122_R_BA37_10 | R | 0.013175647 |
| | | | | 126_No_BA_region | - | 0.028390507 |
| | | | | 140_L_BA10_1 | L | 0.011781666 |
| | | | | 142_No_BA_region | - | -0.012024517 |
| | | | | 152_L_BA23_2 | L | 0.010772799 |
| | | | | 159_No_BA_region | - | -0.008207227 |
| | | | | 193_L_BA10_2 | L | 0.023553666 |
| | | | | 197_L_BA6_3 | L | 0.015629508 |
| | | | | 203_L_BA6_6 | L | 0.012659086 |
| | | | | 26_R_BA20_2 | R | -0.021814024 |
| | | | | 265_L_BA38_3 | L | -0.010083499 |
| | | | | 273_L_BA48_2 | L | 0.011987059 |
| | | | | 47_R_BA23_1 | R | 0.010771693 |
| | | | | 6_R_BA10_4 | R | 0.021289696 |
| | | | | 70_R_BA38_1 | R | -0.056191491 |
| | | | | 75_R_BA39_4 | R | 0.013492614 |
| | | | | 89_R_BA38_3 | R | -0.011975584 |
| | | | | 98_R_BA49_2 | R | -0.020715617 |
| GMV | | | | | | |
| | **216_L_BA13_2** | **L** | **0.241651** | 165_L_BA18_8 | L | -0.381766958 |
| | 37_R_BA10_2 | R | -0.22552 | 121_R_BA18_6 | R | 0.276806087 |
| | 65_R_BA11_2 | R | -0.1914 | 238_L_BA21_3 | L | -0.285323298 |
| | | | | 134_R_BA47_2 | R | 0.21385588 |
| | | | | 169_L_BA45_2 | L | -0.267450822 |
| | | | | 170_L_BA11_2 | L | -0.216098831 |
| | | | | 179_L_BA47_1 | L | 0.204884316 |

| | | | | | |
|---|---|---|---|---|---|
| | | | 189_L_BA47_2 | L | 0.216252279 |
| | | | 19_R_BA21_3 | R | 0.276279269 |
| | | | 207_L_BA13_3 | L | 0.242840816 |
| | | | 210_L_BA21_4 | L | 0.257129934 |
| | | | **216_L_BA13_2** | **L** | **0.178065683** |
| | | | 234_L_BA21_2 | L | -0.257763501 |
| | | | 252_L_BA37_6 | L | 0.265176998 |
| | | | 26_R_BA20_2 | R | -0.234165018 |
| Psychometric | | | | | |
| | AUD3_ | 0.07392362 | DMQ_2 | | 0.142307918 |
| | **SUB2a** | **-0.1036098** | DMQ_9 | | -0.121456934 |
| | | | BIS_22 | | -0.118896235 |
| | | | edu2 | | 0.110014228 |
| | | | **SUB2a** | | **-0.096338165** |
| | | | BIS_24 | | -0.101404791 |
| | | | DRSEQ_12 | | 0.104640749 |
| | | | BDI20 | | 0.094598532 |

*Note: **bold** is significant for predicting both baseline as well as follow-up AUDIT. **<u>bold + underlined</u>** is in different direction*

Figure S1: Flow chart of the entire data collection

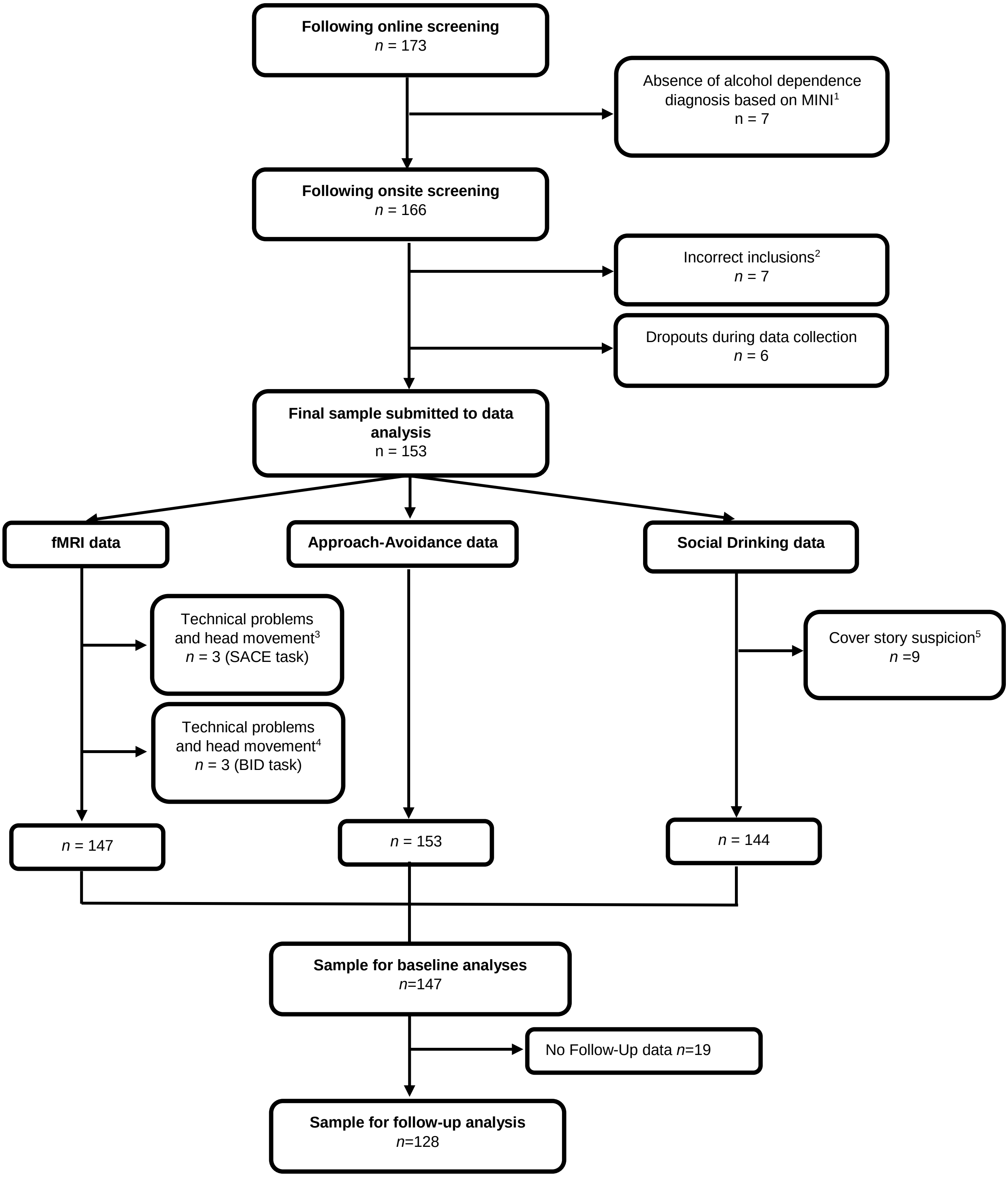


*[1]In the context of the larger project, a severe group of "dependent drinkers" was defined based on AUDIT scores, weekly drinking scores and DSM criteria for alcohol dependence using a MINI interview. Some participants who showed signs of problematic and heavy alcohol use during online screening did not meet the criteria for alcohol dependence following the onsite screening. We decided to exclude these participants, as we deemed that this inconsistency between self-report and DSM-based measures made their categorization uncertain.[2] Incorrect inclusions did not meet the combined requirement for the AUDIT and weekly drink score to be included in the "light drinkers" or " at-risk drinkers" group, also in the context of the larger project. The data from these 7 participants were discarded before performing any data analysis.[3] For the Social-Alcohol Cue Exposure task*

*(SACE), data from one participant was missing, because we were unable to correct for his impaired vision and data from two participants were discarded because they did not pay attention during the task as they did not respond to the control pictures using a button-press. [4]For the Beer-Incentive Delay task (BID), data from one participant was missing because we had technical problems with the pumps to deliver beer and water, and data from two participants were discarded because there was too much head movement. [5]Nine individuals appeared to be aware of the true purpose of the Bar-Lab sessions as they reported on the 'social influence' as an aim of the study.*

Supplementary information on task descriptions and preprocessing details.

*Social Alcohol Cue-Exposure task (SACE) (as published in (5))*

We used a modified version of a passive viewing Cue Exposure task (12), including four conditions of interest (SA: social alcohol, SS: social soda, NA: non-social alcohol, NS: non-social soda, similar to (13)), and one control condition (animal pictures) to which participants had to respond by a button press to ensure they paid attention to the cues. The non-social cues were pictures of beer or soda bottles without any human beings present, while the social pictures showed two or more male and/or female individuals drinking beer or soda while interacting with each other in a social setting, such as a bar or at home. Alcohol and soda pictures were matched one-on-one in terms of social setting, and the number and gender of people present. Twenty cues for each condition were presented in a block design: there were four epochs each consisting of four blocks with five consecutively presented pictures of the same condition (SA, SS, NA or NS). Each picture was presented once for 4.8 s. Blocks were presented in a randomized order which was the same for all participants. There was a 6 s delay (fixation cross) in between each block. Between the four epochs, participants had a 16 s break (fixation cross). Each epoch included one control cue—an animal picture presented for 4.8 s to which participants had to respond—either at the beginning or at the end of a block (except for the last epoch in which two control cues were presented). Total task duration (including 10 practice trials) was approximately 10 min. The main outcome measure of this task was brain cue-reactivity to social alcohol pictures (versus social soda pictures) compared with non-social alcohol pictures (versus non-social soda pictures), that is, the interaction contrast ((SA>SS)-(NA>NS)).

*Approach bias task (as published in (5))*

After scanning, approach biases were measured outside of the scanner using a well-validated stimulus-response compatibility (SRC) task (13, 14). Participants were presented with the exact same pictures as in the SACE task (i.e., SA, SS, NA and NS) and were instructed to either approach or avoid each picture, based on the alcohol content of the picture: "Approach Alcohol" (and "Avoid Soda") or "Avoid Alcohol" (and "Approach Soda"). Every picture was presented for 2,000 ms with a manikin randomly positioned above or below the picture. Participants could approach or avoid the picture by pressing the "up" or "down" keyboard button and thereby moving the manikin in the corresponding direction. After incorrect responses, a red cross was shown for 2,000 ms, and after omissions, "please respond faster" was also shown for 2,000 ms. Participants completed four blocks of 32 trials each: two blocks with only social pictures (one with an "Approach Alcohol" instruction, and one with an "Avoid Alcohol" instruction) and two blocks with only non-social pictures (one with an "Approach Alcohol" instruction and one with an "Avoid Alcohol" instruction). All pictures were presented twice, once within the "Approach Alcohol" block and once within the "Avoid Alcohol" block. The order of task blocks was counterbalanced across participants, with the restriction that those who started with a social block completed both social blocks before proceeding to the non-social blocks, and vice versa. Total task dura ion was approximately 10 min. The task was preceded by 16 practice trials in which participants were instructed to approach bird pictures and avoid pictures of other animals. The outcome measure of the SRC task is the approach bias in each of the four conditions (SA, NA, SS and NS). For each condition, this approach bias was calculated by subtracting the mean reaction time observed for the "Approach" instruction from the mean reaction time observed for the "Avoid" condition including successful trials only. Errors, omissions and outliers (responses < 200 ms and > 2,000 ms or 3 SD above the individual mean) were discarded from these calculations (14, 15). Additionally, we calculated an interaction score by subtracting the non-social alcohol bias from the social alcohol bias (i.e., (SA-bias>SS-bias)-(NA-bias>NS-bias)).

*Beer-Incentive Delay task (as published in (4))*

We used a modified version of the monetary incentive delay task (16, 17) in which the rewards were 3-mL sips of either chilled beer or water. The sips were delivered using two StepDos 03RC fluid pumps with tubes that were placed in the participants' mouth. In the anticipation phase, participants first saw a cue informing them about the opportunity to earn beer (yellow triangle) or water (blue square) followed by a variable delay materialized by a fixation cross. Then, a visual target appeared, and participants were instructed to respond to it as fast as possible using a button press, both in the beer and water conditions. If the response was fast enough, positive feedback was provided in the form of a green tick (outcome notification phase), followed by the drink delivery in the mouth, and then swallowing (delivery phase). When the response was too slow, a red cross was presented, ending that trial. Both the beer and water conditions consisted of 30 pseudo-randomized trials. Ten practice trials preceded the task. Reaction times during the practice trials were used to tailor task difficulty to each individual (by adjusting the time limit for responding to the target), which was further continuously adjusted online to ensure an overall success rate of approximately 66% in each condition (17). The task duration was approximately 20 minutes.

*Drinking scores in the Barlab (as published in (5))*
The Bar-Lab was designed to look like a real bar, increasing the ecological validity of the (imitation of ) drinking measures (18). To cover the real aim of the study, participants were told that they took part in a study on the evaluation of alcohol advertisements. Participants were fully debriefed after study completion. During both Bar-Lab sessions, a confederate was present, acting as a participant to facilitate imitation of drinking. Confederates were 20 males aged between 18 and 25, similar to the participants. After entering the Bar-Lab, the participant and confederate were instructed to fill in online questionnaires on demographics, drinking habits and drinking motives, followed by the rating of several non-alcoholic video advertisements in terms of attractiveness. Then, they were asked to sit at the bar, where peanuts and drinks were available, for a break lasting 30 min, before they had to rate video advertisements again. The experimenter offered a drink to the confederate first to set the norm and enable the examination of whether the participant would choose the same drink. Various soda drinks (200 ml) and two types of local beers (250 ml, 5%–5.2% alcohol) were offered. After providing the first drink, the experimenter left the Bar-Lab after explaining to the participant and confederate that they were allowed to get more drinks if they wanted to. Just before the sessions, the confederates were told to either drink one alcoholic beer followed by one soda (hereafter referred to as the "light" condition) or three alcoholic beers (hereafter referred to as the "heavy" condition) during a 30-min "break." Importantly, the confederate was instructed to initiate the drinks, by informing the participant on what he was drinking, and asking the participant if he would like something to drink as well in a neutral tone. Video and audio recordings were made during the sessions to record the number of drinks consumed. Following the break, the participant and confederate were asked again to rate the alcohol advertisements. They were also asked how they felt during the experiment and what they thought the aim of the study was (suspicion check), as well as how they subjectively evaluated the confederate. These evaluations of the confederates were not correlated with the number of alcoholic drinks consumed by the participants, and across all confederates, similar drinking patterns were found. Each participant completed a "light" and a "heavy" session with two different confederates. Session order was counterbalanced across participants and confederates. This procedure allowed us to quantify both imitation of drinking and social drinking in an ecological setting. Imitation scores were calculated by computing the difference in the number of beers consumed by the participant versus the confederate in each session, and then summing the absolute values of these differences. Social drinking scores were calculated by summing the number of beers consumed across both sessions.

*Preprocessing (as published in (5))*
Imaging was conducted on a PRISMA(Fit) 3T Siemens scanner, using a 32-channel head coil. Blood oxygen level-dependent (BOLD) sensitive functional images were acquired with a whole-brain T2*-weighted sequence using multi-echo echoplanar imaging (EPI) (35 axial slices, matrix 64 × 64, voxel size = 3.5 × 3.5 × 3.0 mm, repetition time = 2,250 ms, echo times = [9.4 18.8 28.2 37.6 ms], flip angle = 90°). The BOLD data acquisition sequence was updated during the course of the study, due to the discovery of MRI noise artifacts. The sequence parameters remained identical, except for the slice order which changed from ascending to interleaved. We took some actions in our analyses to (a) remove the artifacts and (b) model the change in scanning sequence halfway through the study (see below). A high-resolution T1 scan was acquired in each participant (192 sagittal slices, field of view 256 mm, voxel size = 1.0 × 1.0 × 1.0 mm, repetition time = 300 ms, echo time 3.03 ms). Pre-processing steps were conducted in SPM8 (www. fil.ion.ucl.ac.uk/spm). For each volume, the four echo images were combined into a single one, weighing all echoes equally. Standard pre-processing steps were performed on the functional data: realignment to the first image of the time series, co-

registration to the structural image, normalization to MNI space based on the segmentation and normalization of the structural image, and spatial smoothing with an 8-mm Gaussian kernel. In addition, two cleaning methods were incorporated into the pipeline to ensure optimal removal of artifacts and thorough de-noising of the data: (a) a principal component analysis (PCA) to filter out slice-specific noise components (19) before pre-processing and (b) an independent component analysis (ICA)-based automatic removal of motion artifacts using FSL (http://www.fmrib.ox.ac.uk/fsl) after pre-processing (ICA-AROMA; (20, 21)). This pipeline has previously been found to be efficient to take care of the MRI noise artifacts identified in the first half of our data (22).
